# Age-Adjusted Cerebellar Volumetry and Developmental Outcomes in Extremely Low Birth Weight Infants

**DOI:** 10.64898/2026.09.29.755417

**Authors:** Lin Yao, Amanda M. Brown, Junseok Park, Steven Lazar, Stephen Kralik, Christopher J. Rhee, Roy V. Sillitoe, Jason S. Gill

## Abstract

**Background:** Infants with extremely low birth weight (ELBW) are at high risk for neurodevelopmental impairment, but the relationship between cerebellar structure and later developmental outcome remains incompletely defined.

**Methods:** In this retrospective single-center study, we examined total cerebellar volume on clinically acquired brain magnetic resonance imaging (MRI) in ELBW infants while accounting for postmenstrual age (PMA) at imaging.

**Results:** Of 65 infants screened, 42 were included. Mean gestational age at birth was 25.54 ***±*** 1.68 weeks, mean birth weight was 733.2 ***±*** 118.7 g, and mean PMA at MRI was 43.02 ***±*** 6.53 weeks (range, 31.57–68.00). Mean cerebellar volume was 22.17 ***±*** 9.63 cm^**3**^. Independently adjudicated developmental outcomes comprised normal development in 16 infants, isolated or specific impairment in 12, and moderate/severe impairment in 14. Unadjusted cerebellar volumes did not differ across developmental groups (P = 0.490) or qualitative MRI categories (P = 0.875). However, the PMA-by-developmental-category interaction was significant (P = 0.002), with estimated cross-sectional slopes of 1.72, 1.56, and 0.39 cm^**3**^ per week, respectively. The interaction remained significant within 36–46 weeks PMA (n = 29; P = 0.011) and after adjustment for small-for-gestational-age status. Sensitivity analyses indicated uncertainty in the stability of the interaction.

**Conclusions:** These findings suggest that age-informed cerebellar volumetric patterns may provide developmental information not evident from average volumes alone and support prospective validation with standardized imaging and developmental assessments. As a result, cerebellar volumes obtained early in life may hold prognostic information for ELBW infants.

## 1 Introduction

Infants with extremely low birth weight (ELBW; <1000 g) remain at high risk for adverse neurodevelopmental outcomes despite substantial improvements in survival over recent decades [1, 2]. Although ELBW is defined by birth weight rather than gestational age, it substantially overlaps with extreme prematurity; a large multicenter ELBW cohort reported a mean gestational age of approximately 26 weeks, illustrating the substantial overlap between these high-risk populations [3]. Cognitive, language, behavioral, and motor impairments remain common among survivors, with the greatest burden generally observed among infants born at the earliest gestational ages [4–6]. Although cerebral palsy represents an important component of this morbidity, many survivors experience cognitive, language, behavioral, or executive-function difficulties without severe motor impairment [7–9]. These patterns underscore that adverse neurodevelopmental outcomes after preterm birth reflect not only focal destructive brain injury but also disturbances of brain maturation during a period of rapid development. The cerebellum has emerged as an important contributor to neurodevelopment following preterm birth [10, 11]. Beyond its established role in motor control, the cerebellum contributes to cognition, language, executive function, and behavioral regulation through extensive reciprocal connections with cerebral cortical regions [12, 13]. These functions overlap with the cognitive, language, behavioral, and motor impairments observed in children born preterm, providing a strong biologic rationale for examining cerebellar development in relation to later neurodevelopmental outcomes.

The vulnerability of the cerebellum to preterm birth is closely related to its rapid development during late gestation. Cerebellar volume increases approximately four-to fivefold between 24 and 40 weeks’ gestation, with growth that exceeds that of the cerebrum during the same period [11, 14, 15]. The surface area of the cerebellar cortex expands more than 30-fold during this interval, accompanied by rapid proliferation of granule cell precursors in the external granular layer and their subsequent migration to the internal granule cell layer [11, 16, 17]. This period of accelerated growth, cellular proliferation, differentiation, and migration may render the developing cerebellum
particularly susceptible to disruption following preterm birth.

Cerebellar development may be affected by both direct and indirect consequences of preterm birth, including cerebellar hemorrhage, supratentorial white-matter injury, systemic illness, and exposure to selected postnatal therapies or neurotropic medications [11, 18–22]. Supratentorial brain injury may also influence cerebellar development through disruption of cerebro-ponto-cerebellar trophic signaling [23, 24]. Together, these mechanisms provide a biologic rationale for examining quantitative cerebellar development in relation to later neurodevelopmental outcomes in preterm infants.

Quantitative MRI studies have demonstrated smaller cerebellar volumes in preterm infants compared with term-born reference populations. Srinivasan and colleagues reported lower cerebellar volumes in very preterm infants at term-equivalent age compared with term-born controls (25.4 cm^3^ vs 26.9 cm^3^), with smaller volumes also associated with supratentorial lesions [25]. Subsequent studies have shown that reduced cerebellar volume may also occur in preterm infants without visible cerebellar injury, suggesting that altered cerebellar development is not limited to infants with overt focal cerebellar lesions [26, 27].

Quantitative neuroimaging may provide prognostic information beyond categorical lesion scoring. In preterm infants, quantitative measures of white-matter injury have been associated with later motor, cognitive, and language outcomes [28]. However, quantitative approaches evaluating total cerebellar volume in relation to later developmental outcomes remain comparatively less developed [25–28].

Studies evaluating cerebellar pathology in preterm infants have often focused on cerebellar hemorrhage as a categorical marker rather than on quantitative cerebellar volume. Hemorrhage classification identifies infants with overt structural injury but may not capture the broader spectrum of altered cerebellar development associated with preterm birth [29, 30]. Reduced cerebellar volume can occur in the absence of visible cerebellar hemorrhage or other focal cerebellar injury, supporting the possibility that quantitative volumetry may detect developmental differences not apparent on qualitative lesion assessment [25–27, 29, 30]. Quantitative cerebellar volumetry may therefore complement categorical injury assessment by providing a continuous measure of cerebellar structure.

Longitudinal studies support the clinical relevance of cerebellar development in preterm infants. Matthews and colleagues found that cerebellar volume around term-equivalent age and subsequent cerebellar development were associated with later cognitive, language, and motor outcomes in very preterm children [31]. More recently, Rapos and colleagues examined cerebellar subregions at 8 years of age and found that neonatal clinical exposures, including mechanical ventilation, invasive procedures, and opioid exposure, were associated with reduced cerebellar volumes years later, with smaller volumes relating to poorer cognitive and motor outcomes [32]. Together, these studies support a relationship between cerebellar development and later neurodevelopment, while also highlighting heterogeneity in imaging timing, developmental assessment, and follow-up across the existing literature. An outstanding question remains regarding whether within-group cerebellar volume derived from MRIs of ELBW infants obtained in the NICU can inform developmental outcomes. Such a finding would hold enormous value for prognostication of developmental outcomes in premature infants.

A critical gap remains regarding how total cerebellar volume measured on clinically obtained MRI relates to later developmental outcomes in ELBW infants, particularly when age at imaging varies. Neurodevelopmental impairments may emerge or become more apparent as children age and developmental demands increase, while follow-up timing and outcome measures vary substantially across existing studies [33, 34]. The relationship between cerebellar volume, age at MRI, and later clinically documented developmental outcome in a racially and ethnically diverse ELBW cohort therefore remains incompletely understood.

Clinically obtained brain MRI during early infancy provides an opportunity to characterize brain injury and maturation in high-risk preterm infants across a range of postmenstrual ages. Quantitative cerebellar volumetry derived from these clinically acquired studies may offer complementary information about cerebellar development beyond qualitative structural assessment. Establishing whether such quantitative measures are associated with later developmental outcomes is an important step toward determining their potential role in longitudinal developmental assessment and risk stratification.

Accordingly, we conducted a retrospective cohort study of ELBW infants who underwent clinically obtained brain MRI suitable for quantitative cerebellar volumetry. The primary objective was to evaluate the relationship between total cerebellar volume and later developmental outcome using a clinically adjudicated hierarchical classification of normal development, isolated or specific developmental impairment, and moderate/severe neurodevelopmental impairment. Because cerebellar volume varies substantially with age during this developmental period, we specifically examined whether the cross-sectional relationship between cerebellar volume and age at MRI differed across developmental outcome categories. Secondary objectives were to characterize qualitative MRI findings and to assess whether the observed developmental associations remained evident after restriction to a 36–46-week MRI-age window and after accounting for small-for-gestational-age status.

## 2. Methods

### 2.1 Study Design and Population

We conducted a retrospective single-center cohort study of extremely low birth weight (ELBW) infants born at Texas Children’s Hospital between January 2016 and January 2022. ELBW was defined as birth weight <1000 g. The study sample was drawn from a larger parent cohort established under a separate protocol approved by the Baylor College of Medicine Institutional Review Board (IRB). This parent cohort also served as the source population for the study by Chao et al. [35]. Eligibility for the parent cohort required premature birth, a birth weight of 401–1000 g, and an umbilical artery catheter (UAC) placed for clinical reasons. Exclusion criteria included major complex congenital anomalies, hydrops fetalis, poor skin integrity, or being in extremis.

The present study was approved by the Baylor College of Medicine IRB (H-55299), with a waiver of informed consent. Within the parent cohort, 65 ELBW infants born during the study period were identified and screened for inclusion. Eligibility required an available brain MRI suitable for reliable quantitative cerebellar volumetry. Exclusions were based on MRI unavailability, inadequate image quality, or imaging otherwise unsuitable for reliable volumetric analysis. The final analytic cohort comprised 42 infants.

### 2.2 Data Collection and Clinical Variables

Demographic, perinatal, and developmental data were extracted from the electronic health record using an institutional, HIPAA-compliant large language model (LLM)–assisted data platform. All extracted variables and note-derived developmental information were manually audited and curated against the source clinical records before inclusion in the final analytic dataset.

Gestational age at birth was recorded in weeks and days and converted to decimal weeks as weeks plus days divided by seven. Postmenstrual age (PMA) at MRI was calculated by adding postnatal age in weeks to gestational age at birth. MRI examinations were obtained as part of clinical care, and imaging timing was not standardized across the cohort.

Small-for-gestational-age (SGA) status was determined using sex-specific Fenton 2013 birth-weight-for-gestational-age thresholds. SGA was defined as birth weight below the 10th percentile and severe SGA as birth weight below the 3rd percentile [36]. Thresholds were applied at the completed gestational week.

Developmental outcome classifications were assigned following manual review of the available longitudinal clinical documentation, as described in Section 2.3.

### 2.3 Developmental Outcome Classification

Developmental outcomes were independently classified by two pediatric neurologists (LY and JG), blinded to MRI findings, through manual review of longitudinal clinical documentation. Disagreements were resolved through discussion to reach a consensus classification.

Follow-up documentation was considered adequate when it contained an inpatient or outpatient encounter note with sufficient information on the patient’s developmental status and relevant diagnoses to assign an outcome category. Classification as a normal outcome required affirmative documentation of age-appropriate development rather than the absence of a recorded diagnosis alone. The date of the encounter supporting the final classification was recorded.

A study-specific, clinically adjudicated hierarchy was used, informed by established distinctions between domain-specific and global developmental delay [37], functional classification of cerebral palsy [38], and hierarchical assessment of neurodevelopmental impairment after extremely preterm birth [39]. Infants were assigned to one of three mutually exclusive categories:

1. Normal development: affirmative documentation of age-appropriate development, without a documented developmental disorder or persistent developmental impairment at the latest adequate follow-up.
2. Isolated or specific developmental impairment: documented domain-specific developmental impairment, including speech/language or motor impairment, or cerebral palsy with Gross Motor Function Classification System (GMFCS) level I–II, without qualifying global or major multidomain impairment.
3. Moderate/severe neurodevelopmental impairment: documented global developmental delay, comparable significant multidomain impairment, or cerebral palsy with GMFCS level III–V. Children with autism spectrum disorder were assigned to this category when accompanying developmental impairment met these criteria.

Global developmental delay was distinguished from isolated developmental delay by documented significant involvement of at least two developmental domains, consistent with established clinical definitions [37]. The separation of cerebral palsy with GMFCS levels I–II from levels III–V was informed by the Canadian Neonatal Follow-Up Network classification described by Richter et al. [38]. When criteria for more than one category were present, the highest qualifying category took precedence. Therapy referral, early-intervention enrollment, developmental concern, or an abnormal screening result alone was insufficient to establish developmental impairment for this study, consistent with the distinction between developmental surveillance, screening, and diagnostic evaluation [40]. Because standardized developmental test scores were not uniformly available, these categories represented clinical groupings rather than uniform psychometric severity thresholds.

### 2.4 Qualitative MRI Classification

Clinical brain MRI findings were classified using a prespecified qualitative framework. Final clinical neuroradiology reports were retrospectively reviewed by a single board-certified neuroradiologist (SK), blinded to developmental outcomes.

MRI findings were categorized as normal, nonspecific abnormalities, or parenchymal and/or hemorrhagic brain injury. Nonspecific abnormalities included delayed myelination, mild ventricular prominence, a simplified gyral pattern, undersulcation, or mild white matter volume loss without a discrete focal destructive lesion. Parenchymal and/or hemorrhagic brain injury included intraventricular or cerebellar hemorrhage, focal parenchymal injury, periventricular leukomalacia, hypoxic-ischemic injury, or other focal structural abnormalities.

### 2.5 Cerebellar Volumetry

A board-certified neuroradiologist (SK) measured cerebellar volume for each patient by manually contouring the cerebellum on each applicable 1-mm axial slice of the three-dimensional T1-weighted MRI sequence using a picture archiving and communication system (PACS). Total cerebellar volume was calculated by summing the contoured areas across slices and multiplying by the slice thickness, and was expressed in cubic centimeters (cm^3^).

### 2.6 Statistical Analysis

Continuous variables were summarized as means and standard deviations, with ranges where appropriate, and categorical variables as counts and percentages. Cerebellar volumes were compared across developmental outcome groups using one-way analysis of variance (ANOVA) and Welch’s ANOVA. Associations between cerebellar volume and gestational age at birth, PMA at MRI, and birth weight were assessed using Pearson correlations and simple linear regression. Comparisons across qualitative MRI categories were performed using ANOVA and linear regression adjusted for PMA at MRI.

We modeled cerebellar volume as a function of PMA at MRI, developmental outcome category, and their interaction using linear regression. Developmental outcome category was treated as a categorical variable, with normal development as the reference group, and PMA was centered at the analytic sample mean. An omnibus F test evaluated whether the PMA–volume association differed across developmental outcome groups. Group-specific slopes were estimated with 95% confidence intervals, and pairwise slope comparisons were adjusted using the Holm method.

Sensitivity analyses were performed in nested subsets restricted to MRI examinations obtained at 36–46 weeks and 38–42 weeks PMA, inclusive. All infants in the 38–42-week subset were also included in the 36–46-week subset and the full cohort. Additional analyses adjusted for small-for-gestational-age status, used heteroskedasticity-robust standard errors (HC3), and assessed the influence of individual observations through leave-one-out analyses. A secondary analysis compared normal development with any developmental impairment.

Associations of small-for-gestational-age status with cerebellar volume were examined using Welch’s t test and linear regression adjusted for PMA at MRI. The association between SGA status and moderate/severe neurodevelopmental impairment was assessed using Fisher’s exact test.

Analyses were performed using IBM SPSS Statistics, version 29. All tests were two-sided where applicable, with a nominal significance threshold of P <0.05. The Holm method was used to adjust pairwise slope comparisons; no other adjustments for multiple comparisons were applied. Analyses were considered exploratory.

## 3. Results

### 3.1 Cohort Characteristics and Developmental Outcomes

Initial screening identified 65 ELBW infants born at Texas Children’s Hospital during the study period. Twenty-three infants were excluded because MRI was unavailable, image quality was inadequate, or imaging was unsuitable for reliable cerebellar volumetry, yielding a final analytic cohort of 42 infants (Figure 1).

**Fig 1.**
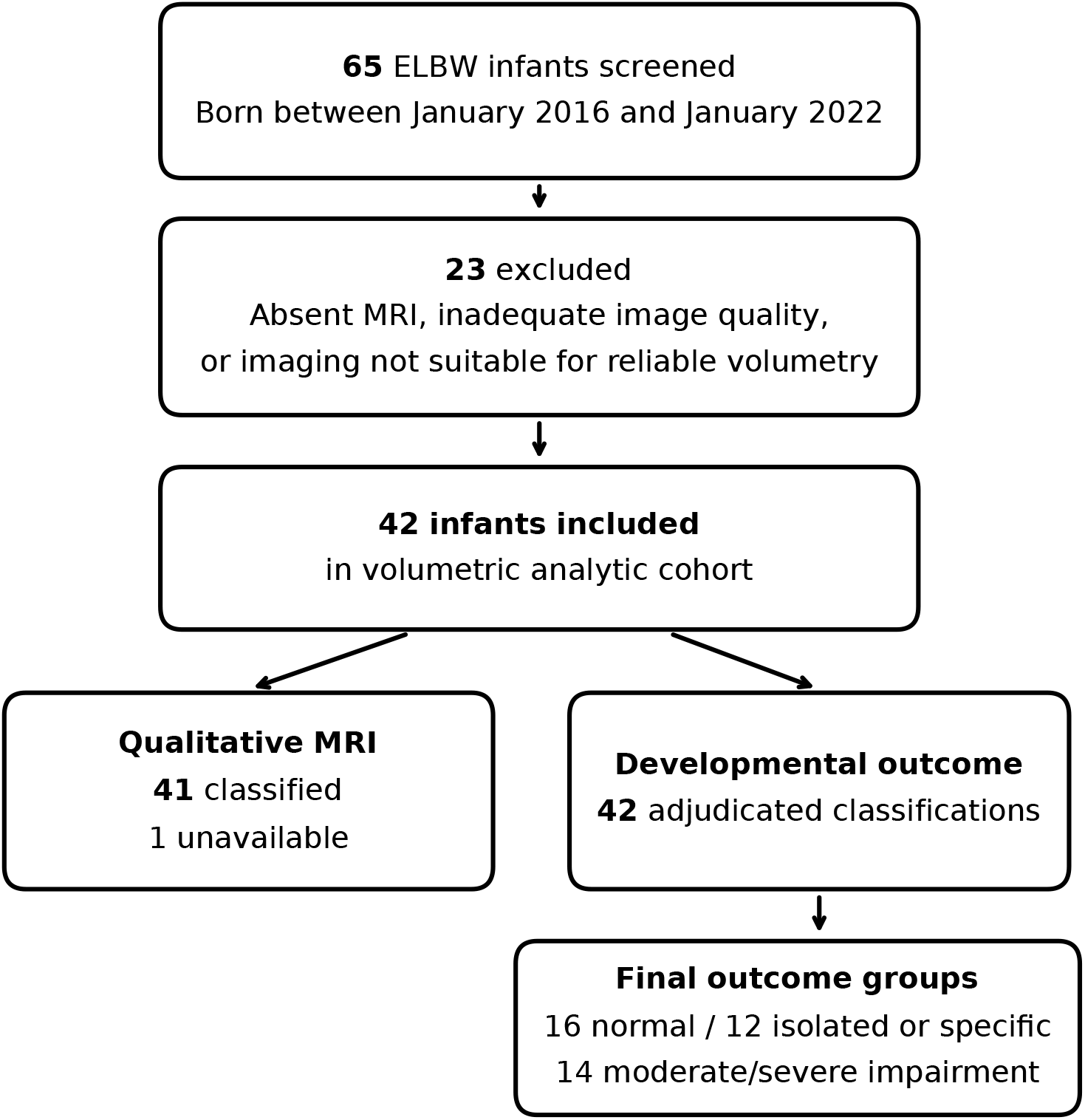
Cohort selection and analytic denominators. All 42 infants had a final adjudicated developmental classification; 41 had evaluable qualitative MRI classification. ELBW, extremely low birth weight

Mean gestational age at birth was 25.54 *±* 1.68 weeks (range, 22 weeks 6 days to 30 weeks 3 days), and 39 infants (92.9%) were born before 28 weeks. Mean birth weight was 733.2 *±* 118.7 g (range, 410–967 g). Twenty infants (47.6%) were female. Recorded race was Black for 23 infants (54.8%), White for 14 (33.3%), Asian for four (9.5%), and mixed race for one (2.4%). Eleven infants (26.2%) were Hispanic. Four infants (9.5%) were classified as SGA, including two (4.8%) with severe SGA (Table 1).

**Table 1.** Cohort characteristics and developmental outcome classification.

| Characteristic | Value |
| --- | --- |
| Final analytic cohort | 42 |
| Gestational age at birth, weeks | $25.54 \pm 1.68$ (22+6 to 30+3) |
| Born before 28 weeks | 39 (92.9%) |
| Birth weight, g | $733.2 \pm 118.7$ (410–967) |
| Female sex | 20 (47.6%) |
| Black race | 23 (54.8%) |
| White race | 14 (33.3%) |
| Asian race | 4 (9.5%) |
| Mixed race | 1 (2.4%) |
| Hispanic ethnicity | 11 (26.2%) |
| PMA at MRI, weeks | $43.02 \pm 6.53$ (31.57–68.00) |
| Cerebellar volume, cm<sup>3</sup> | $22.17 \pm 9.63$ (3.49–44.84) |
| SGA / severe SGA | 4 (9.5%) / 2 (4.8%) |
| Normal development | 16 (38.1%) |
| Isolated or specific impairment | 12 (28.6%) |
| Moderate/severe impairment | 14 (33.3%) |
| Qualitative MRI classification available | 41 (97.6%) |

Values are mean ± SD (range) or n (%). Birth GA range is expressed as weeks+days. Percentages use n = 42. Race and ethnicity were recorded separately; Hispanic ethnicity is not mutually exclusive with race. Severe SGA is included within SGA. PMA, postmenstrual age; SGA, small for gestational age.

All 42 infants received a final developmental outcome classification following clinical adjudication. Sixteen (38.1%) had normal development, 12 (28.6%) had isolated or specific developmental impairment, and 14 (33.3%) had moderate/severe neurodevelopmental impairment.

### 3.2 Qualitative Neuroimaging Profiles

MRI examinations were obtained at a mean PMA of 43.02 *±* 6.53 weeks (range, 31.57–68.00 weeks). Twenty-nine infants underwent MRI at 36–46 weeks PMA, and 14 underwent MRI at 38–42 weeks PMA.

Qualitative MRI classification was available for 41 of 42 infants. Fifteen (36.6%) had normal MRI findings, 14 (34.1%) had nonspecific abnormalities, and 12 (29.3%) had parenchymal and/or hemorrhagic brain injury. The infant without an evaluable qualitative classification remained included in the volumetric and developmental-outcome analyses.

### 3.3 Quantitative Cerebellar Volumetry and Clinical Associations

Mean total cerebellar volume was 22.17 *±* 9.63 cm^3^, with a greater than 12-fold range from 3.49 to 44.84 cm^3^. Mean volumes were 22.02 *±* 5.27 cm^3^ in the normal-MRI group, 23.48 *±* 11.54 cm^3^ in the nonspecific-abnormality group, and 21.61 *±* 11.93 cm^3^ in the parenchymal and/or hemorrhagic injury group. Cerebellar volume did not differ significantly across qualitative MRI categories in unadjusted analysis (P = 0.875) or after adjustment for PMA at MRI (P = 0.719; Figure 2a). Representative MRI images illustrating the range of cerebellar size are shown in Figure 3.

**Fig 2.**
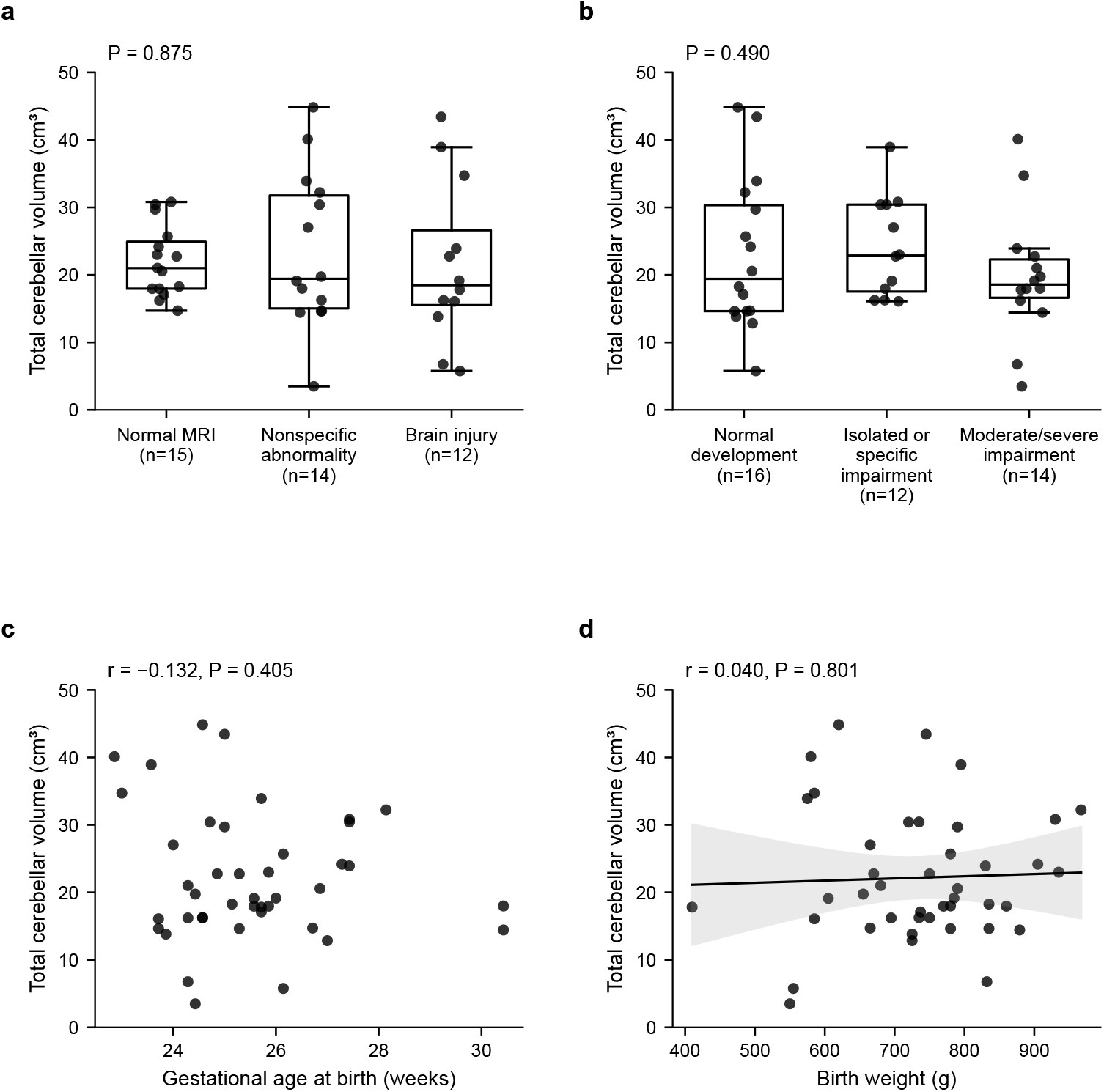
Total cerebellar volume in relation to qualitative MRI findings (a), developmental outcome (b), gestational age at birth (c), and birth weight (d). Boxes show medians and interquartile ranges; whiskers extend to observations within 1.5 interquartile ranges. Points show individual observations, with horizontal jitter in a–b. P values in a–b are from one-way ANOVA; r and P values in c–d are from Pearson correlations using continuous gestational age and birth weight, respectively. The solid line and shaded band in d represent the fitted linear regression and 95% confidence interval. Brain injury denotes parenchymal and/or hemorrhagic brain injury

**fig 3.**
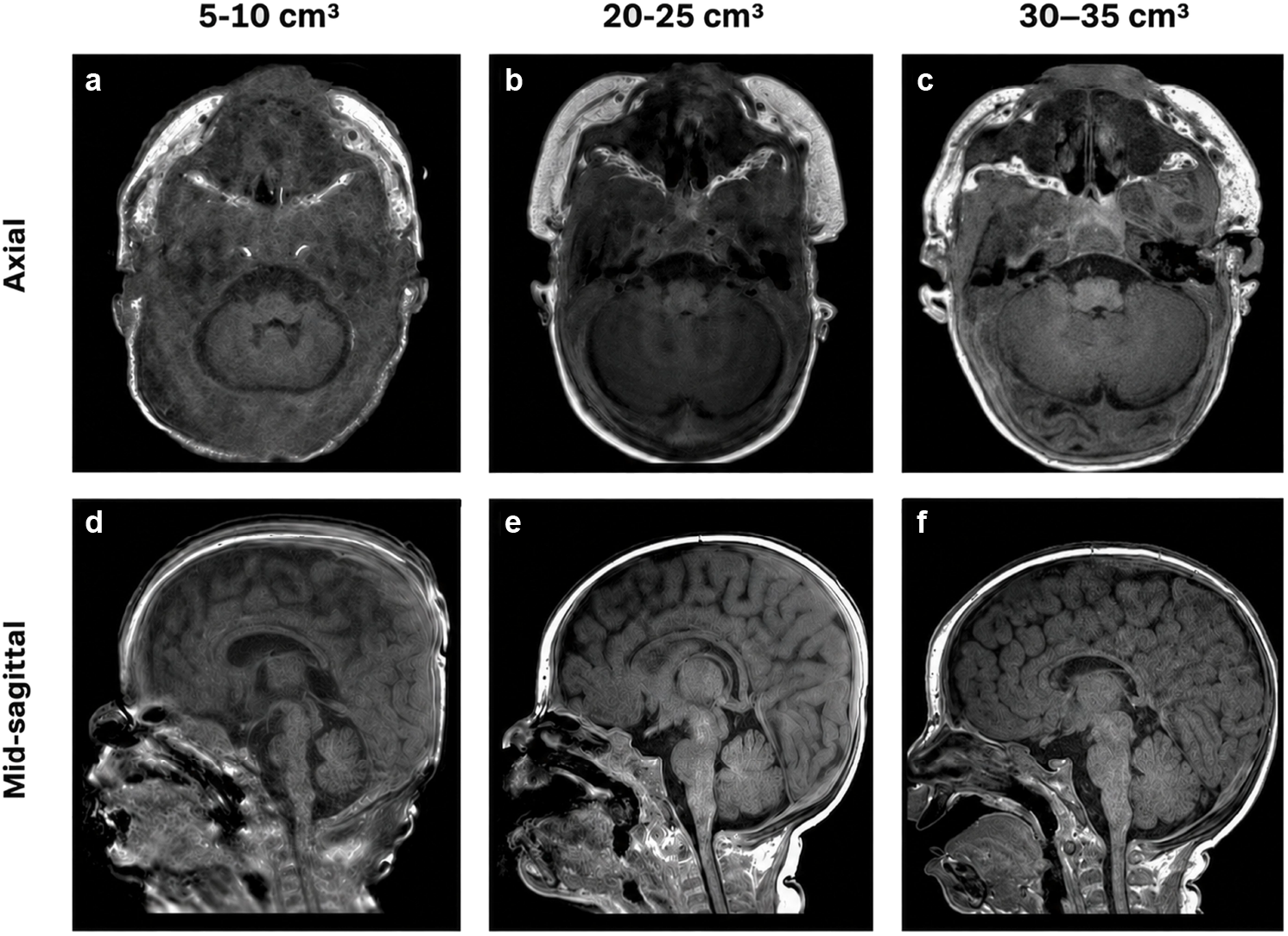
Representative magnetic resonance images from infants with low, intermediate, and higher total cerebellar volumes. axial (a–c) and mid-sagittal (d–f) t1-weighted brain mr images are shown from three representative infants. panels a and d show an infant born at 26 weeks 1 day of gestation with a birth weight of 555 g and a total cerebellar volume of 5.76 cm^3^. panels b and e show an infant born at 25 weeks 6 days of gestation with a birth weight of 935 g and a total cerebellar volume of 22.99 cm^3^. panels c and f show an infant born at 27 weeks 3 days of gestation with a birth weight of 930 g and a total cerebellar volume of 30.81 cm^3^. images are presented for qualitative illustration of the inter-individual variation in cerebellar volume observed in the cohorts

Cerebellar volume was positively associated with PMA at MRI (r = 0.540, P <0.001). In simple linear regression, each additional week of PMA was associated with a 0.80-cm^3^ greater cerebellar volume (95% CI, 0.40–1.19; R^2^ = *™*0.292). Cerebellar volume was not significantly associated with gestational age at birth (r = −0.132, P = 0.405) or birth weight (r = 0.040, P = 0.801; Figure 2c–d).

### 3.4 PMA-Adjusted Cerebellar Volume and Developmental Outcomes

Mean cerebellar volume was 22.89 *±* 11.28 cm^3^ in infants with normal development, 24.08 *±* 7.43 cm^3^ in those with isolated or specific impairment, and 19.72 *±* 9.41 cm^3^ in those with moderate/severe impairment. Unadjusted differences across groups were not statistically significant by one-way ANOVA (P = 0.490) or Welch’s ANOVA (P= 0.430; Figure 2b).

In the primary regression model, the association between PMA and cerebellar volume differed across developmental outcome groups (interaction F (2, 36) = 7.16, P = 0.002; model R^2^ = 0.566). Estimated cross-sectional slopes were 1.72 cm^3^ per week for normal development (95% CI, 1.08–2.35), 1.56 cm^3^ per week for isolated or specific impairment (95% CI, 0.59–2.54), and 0.39 cm^3^ per week for moderate/severe impairment (95% CI, *™*0.05 to 0.83; Table 2 and Figure 4a).

**Table 2.**
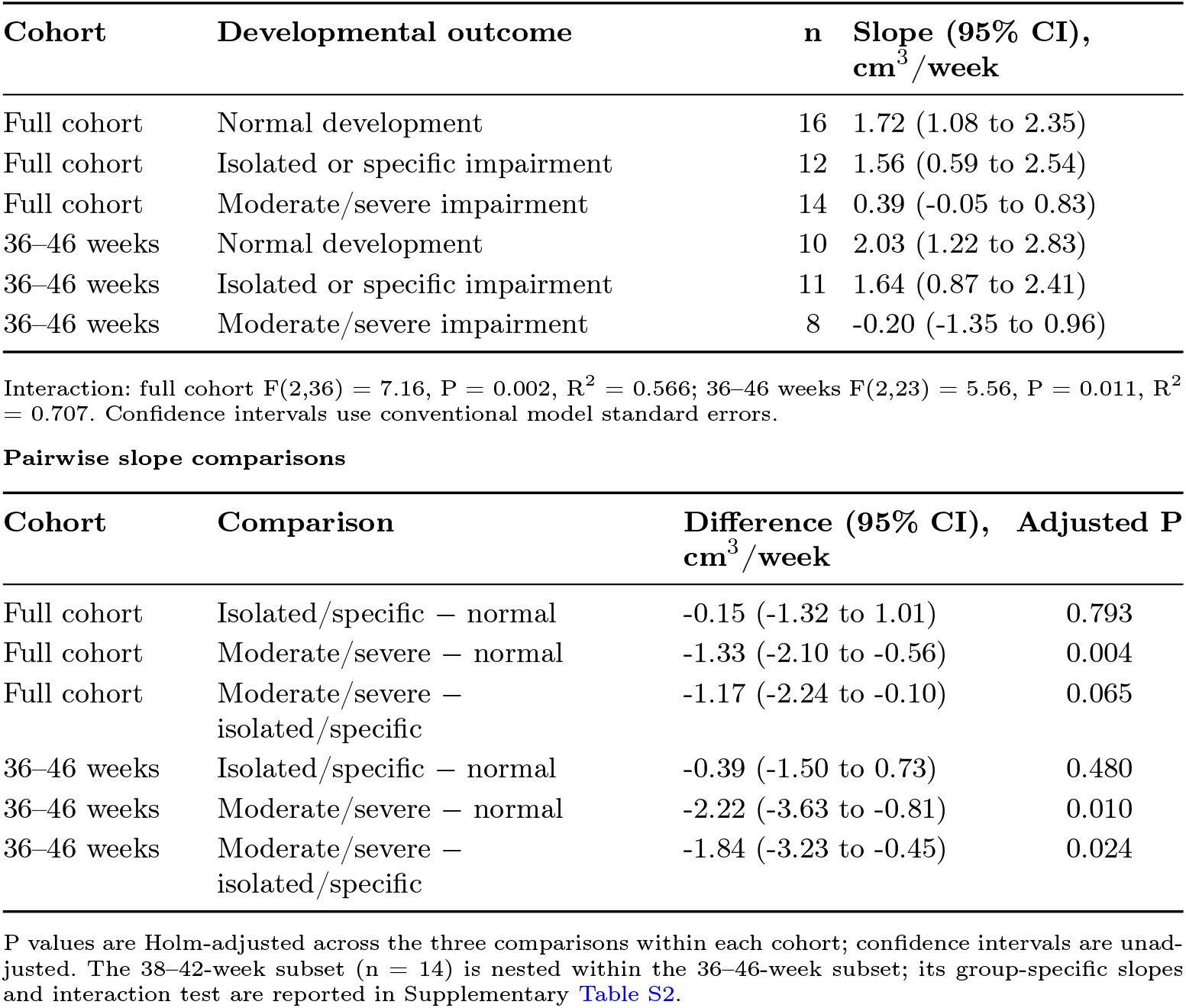
Group-specific cross-sectional PMA–volume slopes.

| Cohort | Developmental outcome | n | Slope (95% CI),<br>cm<sup>3</sup>/week |
| --- | --- | --- | --- |
| Full cohort | Normal development | 16 | 1.72 (1.08 to 2.35) |
| Full cohort | Isolated or specific impairment | 12 | 1.56 (0.59 to 2.54) |
| Full cohort | Moderate/severe impairment | 14 | 0.39 (-0.05 to 0.83) |
| 36–46 weeks | Normal development | 10 | 2.03 (1.22 to 2.83) |
| 36–46 weeks | Isolated or specific impairment | 11 | 1.64 (0.87 to 2.41) |
| 36–46 weeks | Moderate/severe impairment | 8 | -0.20 (-1.35 to 0.96) |

Pairwise slope comparisons
| Cohort | Comparison | Difference (95% CI),<br>cm<sup>3</sup>/week | Adjusted P |
| --- | --- | --- | --- |
| Full cohort | Isolated/specific – normal | -0.15 (-1.32 to 1.01) | 0.793 |
| Full cohort | Moderate/severe – normal | -1.33 (-2.10 to -0.56) | 0.004 |
| Full cohort | Moderate/severe –<br>isolated/specific | -1.17 (-2.24 to -0.10) | 0.065 |
| 36–46 weeks | Isolated/specific – normal | -0.39 (-1.50 to 0.73) | 0.480 |
| 36–46 weeks | Moderate/severe – normal | -2.22 (-3.63 to -0.81) | 0.010 |
| 36–46 weeks | Moderate/severe –<br>isolated/specific | -1.84 (-3.23 to -0.45) | 0.024 |

P values are Holm-adjusted across the three comparisons within each cohort; confidence intervals are unadjusted. The 38–42-week subset ( $n = 14$ ) is nested within the 36–46-week subset; its group-specific slopes and interaction test are reported in Supplementary Table S2.

**Fig 4.**
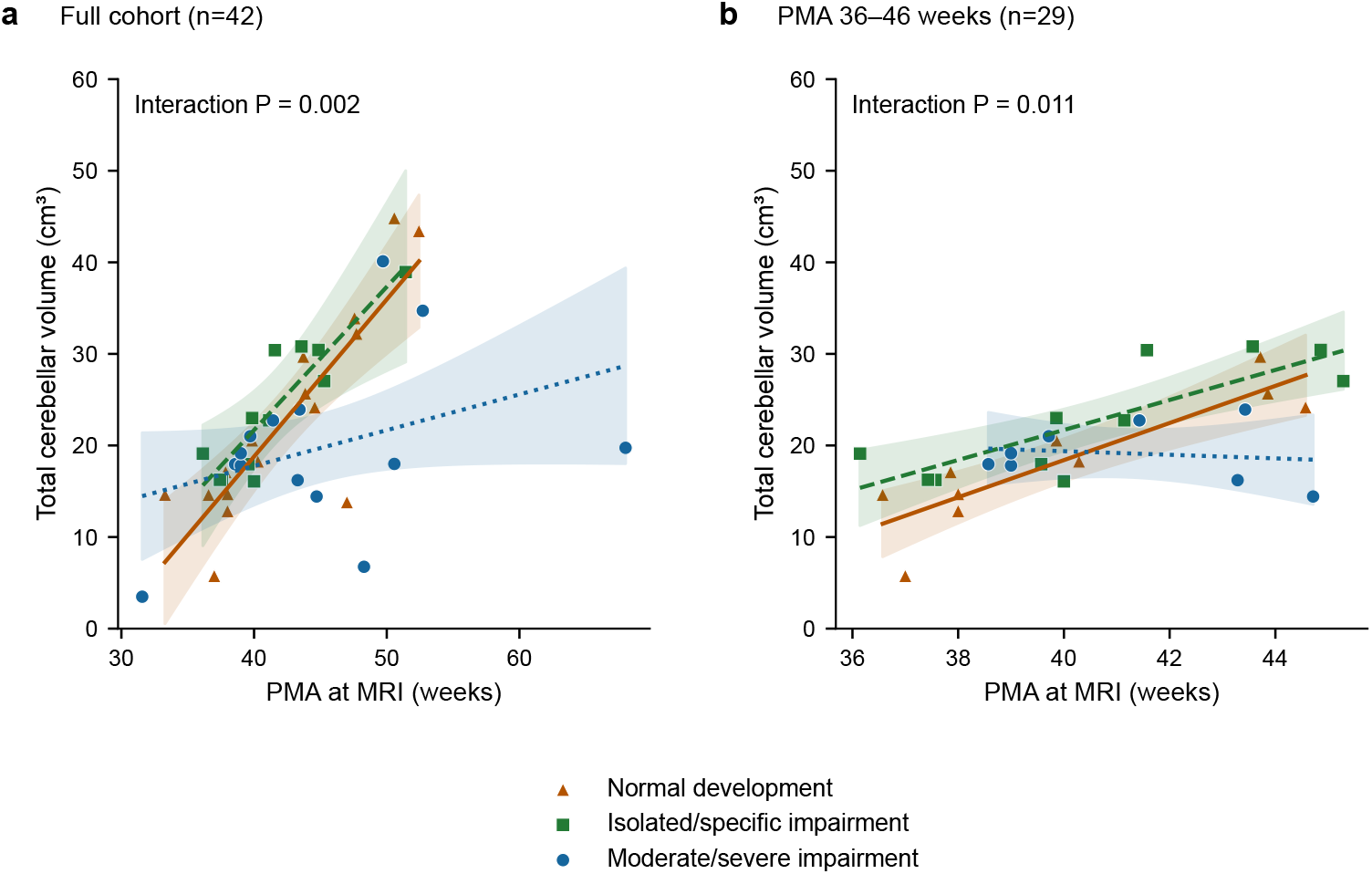
Cross-sectional association between PMA at MRI and total cerebellar volume by developmental outcome in the full cohort (a) and the 36–46-week cohort (b). Lines and shaded bands show fitted group-specific regressions and 95% confidence intervals from the interaction model, limited to each group’s observed PMA range. These slopes represent between-infant associations, not longitudinal growth. PMA, postmenstrual age. Group-specific marker shapes and line styles provide identification in addition to color

The slope was lower in the moderate/severe impairment group than in the normal-development group (difference, *™*1.33 cm^3^ per week; 95% CI, *™*2.10 to 0.56; Holm-adjusted P = 0.004). The comparison between the moderate/severe impairment and isolated or specific impairment groups did not reach significance after Holm adjustment (difference, *™*1.17 cm^3^ per week; 95% CI, *™*2.24 to *™*0.10; adjusted P = 0.065). Slopes did not differ significantly between the isolated or specific impairment and normal-development groups (adjusted P = 0.793). Confidence intervals for pairwise differences were unadjusted.

### 3.5 Sensitivity and Secondary Analyses

The 36–46-week analysis included 29 infants: 10 with normal development, 11 with isolated or specific impairment, and eight with moderate/severe impairment. Unadjusted volumes did not differ significantly across groups (ANOVA P = 0.205; Welch’s ANOVA P = 0.230). The PMA-by-developmental-category interaction remained significant (F(2, 23) = 5.56, P = 0.011; model R^2^ = 0.707).

Within this interval, estimated slopes were 2.03 cm^3^ per week for normal development (95% CI, 1.22–2.83), 1.64 cm^3^ per week for isolated or specific impairment (95% CI, 0.87–2.41), and *™*0.20 cm^3^ per week for moderate/severe impairment (95% CI, *™*1.35 to 0.96). The isolated or specific impairment group did not differ significantly from the normal-development group (adjusted P = 0.480). However, the moderate/severe group had a lower slope than both the normal-development group(Holm-adjusted P = 0.010) and the isolated or specific impairment group (adjusted P = 0.024; Table 2 and Figure 4b).

When outcomes were reduced to normal development versus any developmental impairment, the interaction was significant in the full cohort (P = 0.004) but not within 36–46 weeks (P = 0.101). The unadjusted binary volume comparison was nonsignificant (Welch’s P = 0.727). The narrower 38–42-week analysis included 14 infants—four with normal development, five with isolated or specific impairment, and five with moderate/severe impairment—and did not demonstrate a significant three-level interaction (P = 0.256; Supplementary Table S2).

Sensitivity analyses using robust standard errors and leave-one-out analyses indicated that the interaction was sensitive to model assumptions and individual observations (Supplementary Table S1).

Mean cerebellar volume was lower among SGA infants than among non-SGA infants (14.00 *±* 5.73 versus 23.03 *±* 9.60 cm^3^; Welch’s P = 0.039). After adjustment for PMA, SGA status was associated with an 8.87-cm^3^ lower volume (95% CI, *™*17.22 to *™*0.51; P = 0.038). SGA was present in one of 16 infants with normal development, none of 12 with isolated or specific impairment, and three of 14 with moderate/severe impairment. Its association with moderate/severe impairment was not significant (Fisher’s exact P = 0.100).

Adjustment for SGA retained a significant PMA-by-developmental-category interaction in the full cohort (P = 0.003) and the 36–46-week cohort (P = 0.020).

## 4 Discussion

In this retrospective cohort of 42 extremely low birth weight infants, total cerebellar volume measured on clinically obtained brain MRI demonstrated substantial interindividual variability, ranging more than 12-fold from 3.49 to 44.84 cm^3^. Although mean cerebellar volume did not differ significantly among developmental outcome groups, the relationship between cerebellar volume and PMA at MRI differed according to developmental classification. Estimated age–volume slopes were steeper in infants with normal development and isolated or specific impairment than in those with moderate/severe impairment. The moderate/severe group differed significantly from the normal-development group in the full cohort and from both other groups within the 36–46-week interval. The interaction remained evident after adjustment for SGA status. These findings extend the characterization of cerebellar structure in ELBW infants by showing that its relationship with developmental outcome may become more apparent when imaging age and the extent of developmental impairment are considered together.

A central methodological feature of this study was the use of clinically acquired MRI across a range of PMAs. PMA was positively associated with cerebellar volume, emphasizing the importance of scan timing when interpreting volumetric measurements. Rather than treating this variation solely as a source of confounding, the analysis examined whether the age–volume relationship itself differed across outcome groups. Each infant contributed one MRI examination; therefore, the resulting slopes describe cross-sectional differences between infants rather than within-infant growth rates.

Together, our primary findings suggest that adverse neurodevelopmental outcomes associated with cerebellar hypoplasia in a subset of infants with ELBW may be established by the time these infants leave the NICU, prioritizing future work in establishing whether modifiable risk factors distinguish those with adverse outcomes from those following typical neurodevelopmental trajectories.

### 4.1 Cerebellar volume in ELBW infants

The mean total cerebellar volume in this cohort was 22.17 *±* 9.63 cm^3^. For context, Srinivasan and colleagues reported median term-equivalent cerebellar volumes of 25.4 cm^3^ in very preterm infants and 26.9 cm^3^ in term-born controls [25]. These values provide a descriptive comparison, although differences in imaging age, cohort composition, measurement methods, and summary statistics prevent estimation of a normative deficit. In that study, smaller cerebellar volumes were associated with supratentorial lesions, whereas preterm infants with normal imaging had volumes similar to term-born controls. The present study adds a focused description of volumetric heterogeneity within an exclusively ELBW clinical cohort.

Neither gestational age at birth nor birth weight showed a detectable linear association with cerebellar volume within this ELBW cohort. SGA status, however, was associated with smaller cerebellar volume, including after adjustment for PMA. This distinction suggests that birth weight relative to gestational age may contribute information not captured by absolute birth weight alone, although the SGA analysis involved only four infants.

Together, these findings support evaluating cerebellar volume in relation to both developmental timing and the infant’s clinical context. They also identify questions for subsequent studies: whether prenatal growth, postnatal illness, and treatment exposures account for additional volumetric variation, and whether their associations differ across developmental outcome groups.

### 4.2 Qualitative MRI findings and structural injury

Qualitative MRI review demonstrated substantial structural heterogeneity. Infants were classified as having normal MRI findings, nonspecific abnormalities, or parenchymal and/or hemorrhagic brain injury. The nonspecific group included findings such as delayed myelination, mild ventricular prominence, simplified gyral pattern, undersulcation, and mild white matter volume loss without a discrete focal destructive lesion. The parenchymal and/or hemorrhagic brain injury group included intraventricular or cerebellar hemorrhage, focal parenchymal injury, periventricular leukomalacia, hypoxic-ischemic injury, or other focal structural abnormalities.

Total cerebellar volume did not differ significantly across qualitative MRI categories, either before or after adjustment for PMA. Broad qualitative MRI categories did not distinguish mean cerebellar volume in this cohort. Quantitative volumetry provides a continuous description of structure that is not interchangeable with categorical report-based assessment. Whether this additional characterization improves prediction of developmental outcomes remains a separate question.

Prior studies provide a biological basis for examining injury patterns more specifically. Srinivasan and colleagues linked smaller cerebellar volumes to supratentorial lesions, and Shah and colleagues reported relationships between white matter injury and cerebellar volume [25, 26]. Research on cerebro-cerebellar interactions further suggests that injury in one region may influence development in connected structures [23]. The broad categories used here may combine lesions with differing effects on cerebellar development. Injury-specific analyses incorporating lesion location, extent, and associated supratentorial abnormalities could help reconcile this heterogeneity with the relationships reported in previous studies.

### 4.3 Age-adjusted cerebellar volumetric patterns and developmental outcome

Absolute cerebellar volumes overlapped substantially across developmental outcome groups. Mean volumes were 22.89 cm^3^ in infants with normal development, 24.08 cm^3^ in those with isolated or specific impairment, and 19.72 cm^3^ in those with moderate/severe impairment. The absence of a significant unadjusted difference highlights the limitations of interpreting cerebellar size without considering imaging age and developmental phenotype.

The more informative finding was the interaction between PMA and developmental category. Estimated age–volume slopes were 1.72 cm^3^ per week for normal development, 1.56 cm^3^ per week for isolated or specific impairment, and 0.39 cm^3^ per week for moderate/severe impairment. The moderate/severe group differed significantly from the normal-development group after Holm adjustment. Within the 36–46-week interval, the same overall pattern was observed, with the moderate/severe group differing from both other groups. Adjustment for SGA status retained the interaction, suggesting that SGA alone did not account for the observed pattern.

An important contribution of this study is the distinction between isolated or specific impairment and broader developmental involvement. Within the 36–46-week imaging interval, the PMA-by-outcome interaction remained significant using the three-level classification (P = 0.011), whereas it was not significant when developmental outcomes were reduced to delay versus no delay (P = 0.101). The similar estimated slopes in infants with normal development and isolated or specific impairment, together with the shallower slope in those with moderate/severe impairment, suggest that combining all impairments into a single category may obscure differences associated with broader developmental involvement. These findings highlight the importance of outcome definition when examining relationships between cerebellar structure and development.

This finding complements longitudinal studies linking cerebellar development with later motor, cognitive, and language outcomes. Matthews and colleagues examined cerebellar volume and growth from term-equivalent age to childhood in a broader very preterm population [31]. Our study extends this line of investigation to an exclusively ELBW cohort using clinical MRI and developmental outcomes independently adjudicated from longitudinal clinical records. Its contribution is the identification of an outcome-related age–volume pattern in a clinical setting, providing a specific hypothesis for serial-imaging studies: whether differences in individual cerebellar growth are most pronounced among infants who develop broader neurodevelopmental impairment.

### 4.4 Why total cerebellar volume alone may be insufficient

Total cerebellar volume may not differ significantly across developmental groups because it combines anatomically and functionally distinct regions. Regional specialization for motor and nonmotor functions, including language, executive function, and emotional processing, may therefore be obscured by a single volume measure [12].

Developmental impairment is also heterogeneous within clinically defined categories. Although the present classification distinguishes isolated or specific impairment from global or multidomain involvement, children within each group may have different motor, language, cognitive, and behavioral profiles. More detailed domain-specific assessments may therefore be needed.

Finally, clinical classifications depend on the timing and scope of follow-up. Normal development reflects evidence available at the latest adequate assessment but cannot exclude later-emerging difficulties. The limited correspondence between early testing and later cognitive performance in ELBW cohorts highlights the importance of assessment timing [34]. Standardized evaluations at defined ages, combined with longitudinal documentation, would provide a more complete developmental characterization.

### 4.5 Cohort strengths and generalizability

Several strengths of this study are notable. The cohort focused exclusively on ELBW infants, a group at particularly high risk for neurodevelopmental impairment and cerebellar vulnerability. The study also included a racially and ethnically diverse clinical population, with 54.8% Black infants and 26.2% Hispanic infants. The inclusion of a racially and ethnically diverse population adds clinically relevant evidence from an urban tertiary neonatal center. Although external validation remains necessary, the present cohort provides a clinically relevant foundation for studying cerebellar volumetry in tertiary neonatal care.

Another strength is the linkage of MRI-derived cerebellar volume with longitudinal developmental documentation. All 42 infants received a final outcome classification following independent review by two pediatric neurologists blinded to MRI findings. This design reflects real-world practice, where prognostic counseling and follow-up planning often rely on clinically acquired imaging and longitudinal clinical documentation rather than protocolized research assessments.

### 4.6 Limitations

This retrospective, single-center study is subject to selection bias because MRI acquisition, timing, and developmental follow-up were determined clinically. The modest sample size limited precision, particularly for group-specific slope estimates. Although the interaction remained significant after restriction of imaging age and adjustment for SGA, sensitivity analyses indicated uncertainty in its stability, supporting confirmation in a larger cohort.

Because each infant underwent only one MRI, individual growth trajectories could not be estimated. The decision to use absolute cerebellar volumes rather than volumes normalized to supratentorial volume may fail to take into account interindividual differences in overall brain size. However, the use of absolute cerebellar volumes avoids confounding effects of independent variables that may pathologically reduce brain size, such as supratentorial hemorrhage or white matter injury of prematurity, which would risk overcorrection and the underestimation of cerebellar volumetric reductions. In place of cerebellar volumes normalized to brain volume, the association between cerebellar volume and birth weight was examined separately as a clinical correlate (Figure 2d). Developmental outcomes were assigned using a consistent clinical framework, but standardized developmental test scores and uniform follow-up intervals were unavailable. Broad injury categories, unmodeled NICU exposures, and the small SGA subgroup further limited interpretation. These limitations underscore the need for larger-scale prospective studies.

### 4.7 Future directions

Future studies should build on the observed age–volume pattern using prospective cohorts with standardized imaging intervals and repeated MRI. Serial measurements would allow direct assessment of whether cerebellar growth differs between normal development, isolated or specific impairment, and moderate/severe impairment. Development of advanced imaging platforms that use causal mediation analysis to understand the reciprocal relationship between supratentorial brain pathology and cerebellar development would be crucial to fully understand how cerebellar volumes are associated with developmental outcomes. Finally, increasing the size of the dataset to increase measurement reproducibility would strengthen the interpretation of regional volumetric findings and allow for granular understanding of the pathology underlying isolated/specific developmental impairments.

Subregional segmentation and standardized domain-specific developmental assessments could further clarify which aspects of cerebellar structure are associated with motor, language, cognitive, and behavioral outcomes. In parallel, incorporating clinical exposures, growth measures, and quantitative injury assessments would help determine whether cerebellar volume adds information beyond established clinical risk factors.

## 5 Conclusion

The present study provides an ELBW-specific foundation for this work. Its principal contribution is the identification of developmental-group differences in the cross-sectional relationship between cerebellar volume and PMA that were not evident from average volumes alone. These findings support continued evaluation of age-informed cerebellar volumetry within a multimodal approach to developmental assessment and future risk stratification.

## Acknowledgements

Facility and overhead support for this study were made possible through Baylor College of Medicine, Texas Children’s Hospital, and the Jan and Dan Duncan Neurological Research Institute. The Texas Children’s Hospital Research and Resources Office was integral in the collection of the primary dataset. The Cerebellum Science Center fostered an environment of collaboration that made the final version of this manuscript possible.

## Declarations

### Author contributions

L.Y. contributed to data curation, formal analysis, visualization, data interpretation, and writing of the original draft. A.M.B. contributed to data curation, analysis, and interpretation. J.P. contributed to data review. S.L. contributed developmental follow-up data and developmental outcome interpretation. S.K. contributed neuroradiologic review, cerebellar volumetric analysis, and imaging interpretation. C.J.R. contributed cohort assembly. R.V.S. contributed to conceptualization and study design. J.S.G. contributed to conceptualization, methodology, cohort design and development, supervision, and data interpretation. All authors contributed to manuscript review and editing and approved the final manuscript.

### Ethics approval and consent to participate

The study was approved by the Baylor College of Medicine Institutional Review Board (H-55299), with a waiver of informed consent. The study was conducted in accordance with the Declaration of Helsinki and its later amendments.

### Data availability

The data supporting this article cannot be made openly available due to HIPAA guidelines. De-identified participant data are available upon reasonable request made to J.S.G. and subject to a data sharing agreement.

### AI Declaration

An institutional, HIPAA-compliant large language model (LLM)-assisted platform was used to support extraction of demographic, perinatal, and developmental information from the electronic health record; all extracted variables and note-derived information were manually audited and curated against the source clinical records before analysis. OpenAI ChatGPT was used during manuscript preparation to assist with LaTeX compilation and reference management. All AI-assisted outputs were reviewed and verified by the authors. The authors retained full responsibility for the study design, statistical analyses, interpretation, citations, and final manuscript content.

## Supplementary material

**Table S1.**
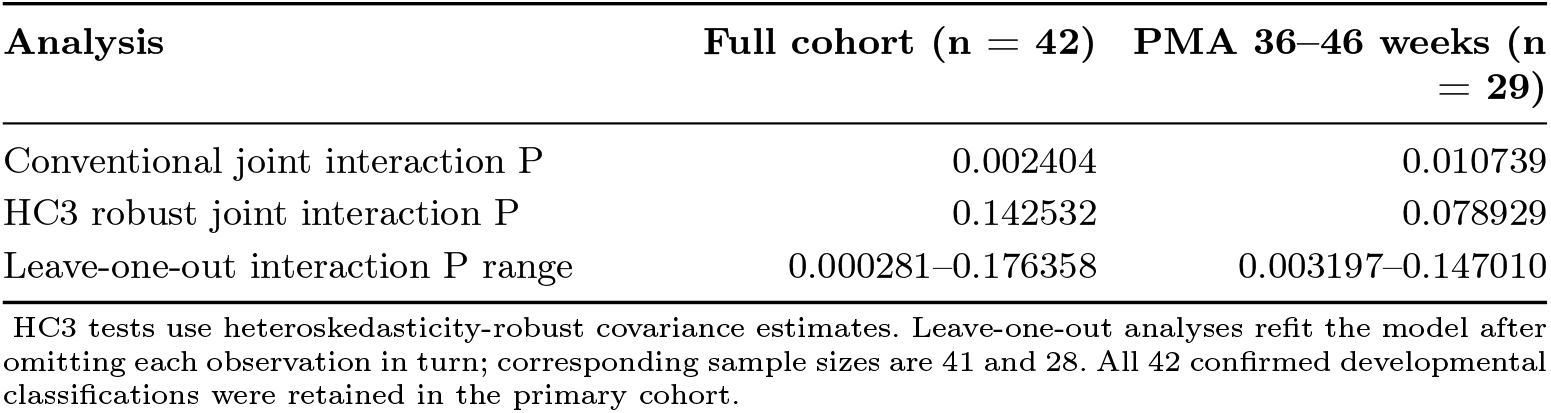
Robustness of the three-level PMA-by-outcome interaction.

**Table S2.**
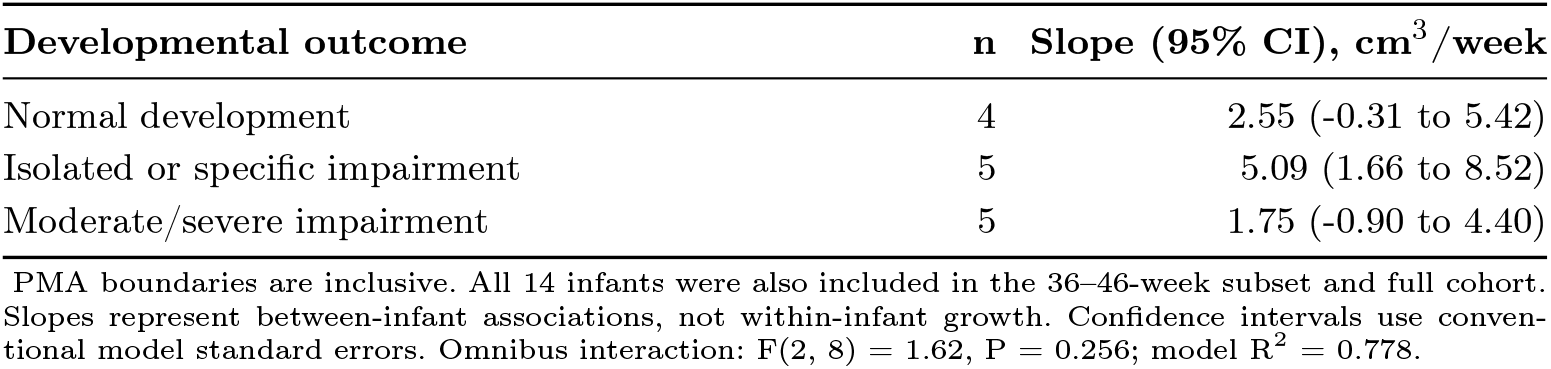
Group-specific cross-sectional PMA–volume slopes in the 38–42-week subset.

