## supplementary tables for "Age-Adjusted Cerebellar Volumetry and Developmental Outcomes in Extremely Low Birth Weight Infants"

### Supplementary material

**Table S1** Robustness of the three-level PMA-by-outcome interaction

| Analysis | Full cohort (n = 42) | PMA 36–46 weeks (n = 29) |
|---|---|---|
| Conventional joint interaction P | 0.002404 | 0.010739 |
| HC3 robust joint interaction P | 0.142532 | 0.078929 |
| Leave-one-out interaction P range | 0.000281–0.176358 | 0.003197–0.147010 |

HC3 tests use heteroskedasticity-robust covariance estimates. Leave-one-out analyses refit the model after omitting each observation in turn; corresponding sample sizes are 41 and 28. All 42 confirmed developmental classifications were retained in the primary cohort.

**Table S2** Group-specific cross-sectional PMA–volume slopes in the 38–42-week subset

| Developmental outcome | n | Slope (95% CI), $cm^3$/week |
|---|---|---|
| Normal development | 4 | 2.55 (-0.31 to 5.42) |
| Isolated or specific impairment | 5 | 5.09 (1.66 to 8.52) |
| Moderate/severe impairment | 5 | 1.75 (-0.90 to 4.40) |

PMA boundaries are inclusive. All 14 infants were also included in the 36–46-week subset and full cohort. Slopes represent between-infant associations, not within-infant growth. Confidence intervals use conventional model standard errors. Omnibus interaction: $F(2, 8) = 1.62$, $P = 0.256$; model $R^2 = 0.778$.
